# SARS-CoV-2 immune complex triggers human monocyte necroptosis

**DOI:** 10.1101/2022.10.13.512127

**Authors:** Leonardo Duarte Santos, Krist Helen Antunes, Gisele Cassão, João Ismael Gonçalves, Bruno Lopes Abbadi, Cristiano Valim Bizarro, Luiz Augusto Basso, Pablo Machado, Ana Paula Duarte de Souza, Bárbara Nery Porto

**Affiliations:** Laboratory of Clinical and Experimental Immunology, Infant Center, School of Life and Health Science, Pontifical Catholic University of Rio Grande do Sul, Porto Alegre, Brazil; Research Center of Functional and Molecular Biology (CPBMF), Pontifical Catholic University of Rio Grande do Sul, Porto Alegre, Brazil; Department of Medical Microbiology and Infectious Diseases, Rady Faculty of Health Sciences, University of Manitoba, Winnipeg, MB, Canada; Biology of Breathing Group, Children’s Hospital Research Institute of Manitoba, Winnipeg, MB, Canada

**Keywords:** SARS-CoV-2, necroptosis, monocyte, Fcγ, receptors, COVD-19 pathogenesis

## Abstract

We analyzed the ability of severe acute respiratory syndrome coronavirus-2 (SARS-CoV-2) itself and SARS-CoV-2-IgG immune complexes to trigger human monocyte necroptosis. SARS-CoV-2 was able to induce monocyte necroptosis dependently of MLKL activation. Necroptosis-associated proteins (RIPK1, RIPK3 and MLKL) were involved in SARS-CoV-2 N1 gene expression in monocytes. SARS-CoV-2 immune complexes promoted monocyte necroptosis in a RIPK3- and MLKL-dependent manner, and Syk tyrosine kinase was necessary for SARS-CoV-2 immune complex-induced monocyte necroptosis, indicating the involvement of Fcγ receptors on necroptosis. Finally, we provide evidence that elevated LDH levels as a marker of lytic cell death are associated with COVID-19 pathogenesis.

## Background

Coronaviruses (CoVs) have been responsible for generating three major outbreaks in recent decades, one caused by middle east respiratory syndrome-CoV (MERS-CoV) and two caused by severe acute respiratory syndromes (SARS-CoV and SARS-CoV-2). Although there are vaccines and treatments available to prevent and control infection by SARS-CoV-2, the mechanisms of COVID-19 pathogenesis are still not well defined. Given the crucial function of monocytes in host defense and their potential activity in the hyper-inflammation seen in COVID-19 patients, understanding the role played by monocytes during COVID-19 is key for tackling the disease’s pathogenic mechanisms [1, 2]. In this regard, the mode of monocyte death may influence COVID-19 outcomes as lytic cell death modes, including necroptosis and pyroptosis, lead to cytokines, chemokines, and alarmins release [3]. Furthermore, necroptotic and pyroptotic cells release lactate dehydrogenase (LDH), which has been widely used as a marker of lytic cell death. Importantly, elevated LDH levels have been used as a correlate for severe COVID-19 [4].

Cytokine-induced monocyte cell death has been suggested to be associated to tissue damage and inflammation during SARS-CoV-2 infection [5]. It has been recently shown that monocytes from COVID-19 patients undergo pyroptosis [6, 7]. Moreover, antibody-mediated SARS-CoV-2 infection of monocytes induces pyroptosis [7]. However, whether SARS-CoV-2 itself and SARS-CoV-2 immune complexes trigger human monocyte necroptosis has yet to be demonstrated.

Here we report that SARS-CoV-2 and SARS-CoV-2-IgG immune complexes promote monocyte necroptosis through RIPK3, MLKL and Syk tyrosine kinase activation. We also show that elevated LDH levels positively correlate with COVID-19 severity.

## Materials and methods

### Reagents

Ham’s F-12 nutrient mix, RPMI 1640 and fetal bovine serum (FBS) were purchased from Gibco (ThermoFischer Scientific, MA, USA). Necrosulfonamide (NSA) was from Millipore (Sigma-Aldrich). Piceatannol was from Cayman Chemical. Histopaque-1077and NEC-1s were purchased from Sigma-Aldrich. GW42X (SYN-1215) from Synkinase was a gift from Dr. Ricardo Weinlich, Hospital Israelita Albert Einstein, Brazil.The GoScript reverse transcription system kit and CytoTox 96 Non-Radioactive Cytotoxicity Assay were from Promega. Brazol reagent was from LGC Biotecnologia. High Capacity cDNA Reverse Transcription Kit was from Applied Biosystems by ThermoFisher Scientific. Anti-spike IgG ELISA was from Euroimmun.

### COVID-19 patients and Ethics statement

Adult patients’ blood was collected from SARS-CoV-2 positive individuals admitted at São Lucas Hospital – PUCRS (Porto Alegre, Brazil), diagnosed by qRT-PCR, from March to July 2021, within 3 weeks of the onset of symptoms and with a maximum of 48 h of hospitalization. Informed consent form was signed before study enrollment. Serum was collected and stored at −20°C. The study was reviewed and approved by the Research Ethics Committee of PUCRS (CEP/PUCRS) under protocol number 30754220.3.0000.5336. All procedures followed the standards established by the Declaration of Helsinki.

### Anti-Spike IgG measurements

Quantitative ELISA was used to determine the concentration of IgG antibodies against the S1 antigen of SARS-CoV-2, following the manufacturer’s instructions. The concentration is expressed as binding antibody units per milliliter (BAU/mL).

### Virus preparation

SARS-CoV-2 was provided by Prof. Dr. Edison Luiz Durigon (ICB-USP). Production of SARS-CoV-2 was performed in Vero cells (CCL81, ATCC) cultured in DMEM F12 medium with 2% FBS at 37°C under 5% CO_2_. To assess viral titer, Vero cells were infected with SARS-CoV-2 in serum-free medium followed by a carboxymethylcellulose plaque assay. The viral titer was determined using TCID_50_. The virus aliquots were stored at −80°C.

### Human monocytes isolation and infection

Peripheral blood mononuclear cells (PBMCs) were isolated from healthy volunteer donors (from both sexes, with a mean age of 30 years). Twenty milliliters of whole blood were collected and subjected to a Histopaque-1077 gradient centrifugation for 30 minutes. PBMCs were seeded at 3 × 10^5^/well in RPMI 1640 medium containing 10% FBS overnight. Afterwards, non-adherent cells were removed by washing, and adherent cells were infected with SARS-CoV-2 (MOI 0.5) for 24 h and LDH release was measured in cell supernatants as described below. To evaluate the role of RIPK1, RIPK3, and MLKL on SARS-CoV-2-induced necrosis, cells were treated with Nec-1s (30 μM), GW42X (2 μM), or NSA (2 μM) for 1 h prior to SARS-CoV-2 infection.

### Preparation of SARS-CoV-2–IgG immune complexes and monocyte stimulation

Serum from COVID-19 patients (COVID-19 serum) and serum obtained from healthy volunteer donors before the pandemic (pre-pandemic serum) were inactivated at 56°C for 30min and then stored at −20°C. SARS-CoV-2 was inactivated under UV light for 30 min and the aliquots were stored at −80°C. To allow immune complexes formation, both COVID-19 and pre-pandemic sera (500 BAU/mL) were incubated with UV-SARS-CoV-2 (MOI 0.5) for 1 h at 37°C. Afterwards, monocytes were incubated with UV-SARS-CoV-2 immune complexes or the mixture of pre-pandemic and UV-SARS-CoV-2 for 24 h at 37°C. LDH release was measured in cell supernatants as described below. To evaluate the role of RIPK1, RIPK3 and MLKL on monocyte necrosis triggered by SARS-CoV-2-IgG immune complexes, cells were treated with Nec-1s (30 μM), GW42X (2 μM) or NSA (2 μM) for 1 h prior to incubation with immune complexes. To determine the role of Syk tyrosine kinase on monocyte necroptosis triggered by UV-SARS-CoV-2 immune complexes, cells were treated with piceatannol (20 μM) for 1 h prior to incubation with the immune complexes.

### LDH release measurements

LDH release was detected in monocyte supernatants using a CytoTox 96 Non-Radioactive Cytotoxicity Assay kit, following the manufacturer’s instructions. Readings were carried out at 490 nm wavelength, using EZ Read 400 microplate reader (Biochrom).

### SARS-CoV-2 N1 gene expression by real-time PCR

Total cell RNA was extracted, and cDNA was synthesized using GoScript reverse transcriptase kit. The expression of the N1 gene was performed using specific primers and probes TaqMan Assay (Applied BioSystems, Thermo Fisher Scientific, Waltham, MA, EUA). The amplification of SARS-CoV-2 nucleocapsid protein gene was performed using the indicated specific primers: forward primer - 5′-GGGGAACTTCTCCTGCTAGAAT-3′, reverse primer - 5′-CAGACATTTTGCTCTCAAGCTG-3′. Quantification of gene expression was carried out using StepOne (Applied Biosystems – ThermoFisher Scientific, MA, USA). A total of 4 ng of cDNA was used for each reaction. The target gene expression was determined using the 2^−ΔΔCt^ method (fold change over control expression).

### Statistical analysis

Data are presented as mean ± SEM. All experiments were performed in triplicates and repeated at least two times. The results obtained were analyzed using GraphPad Prism 8 statistical software package (GraphPad Software, La Jolla, CA, USA). Comparisons between multiple groups were analyzed with one-way ANOVA and a post-hoc Tukey test. When appropriate, a Wilcoxon matched-pairs signed-rank test was employed. The level of significance was set at p ≤ 0.05.

## Results

### SARS-CoV-2 promotes human monocyte necroptosis

To evaluate the effect of SARS-CoV-2 on monocyte death, cells were infected with SARS-CoV-2 (MOI 0.5) for 24 h and cell death was assessed by measuring LDH release on cell supernatants. Our results show that SARS-CoV-2 triggered monocyte necrosis (Fig. 1A). To determine the role of RIPK1, RIPK3 and MLKL on SARS-CoV-2-induced monocyte death, cells were treated with selective inhibitors (NEC-1s, GW-42X, and NSA, respectively) prior to SARS-CoV-2 infection. Inhibiting MLKL almost abolished monocyte death promoted by SARS-CoV-2 (Fig. 1A); however, pharmacological suppression of RIPK1 or RIPK3 did not affect cell death induced by the virus (Fig. 1A), suggesting that SARS-CoV-2 triggers monocyte necroptosis through the activation of MLKL. Next, we sought to understand the role of necroptosis-associated proteins on SARS-CoV-2 N1 gene expression in monocytes. We found that blocking RIPK1, RIPK3 or MLKL significantly reduced the expression of SARS-CoV-2 N1 gene in monocytes (Fig. 1B).

**Fig.1.**
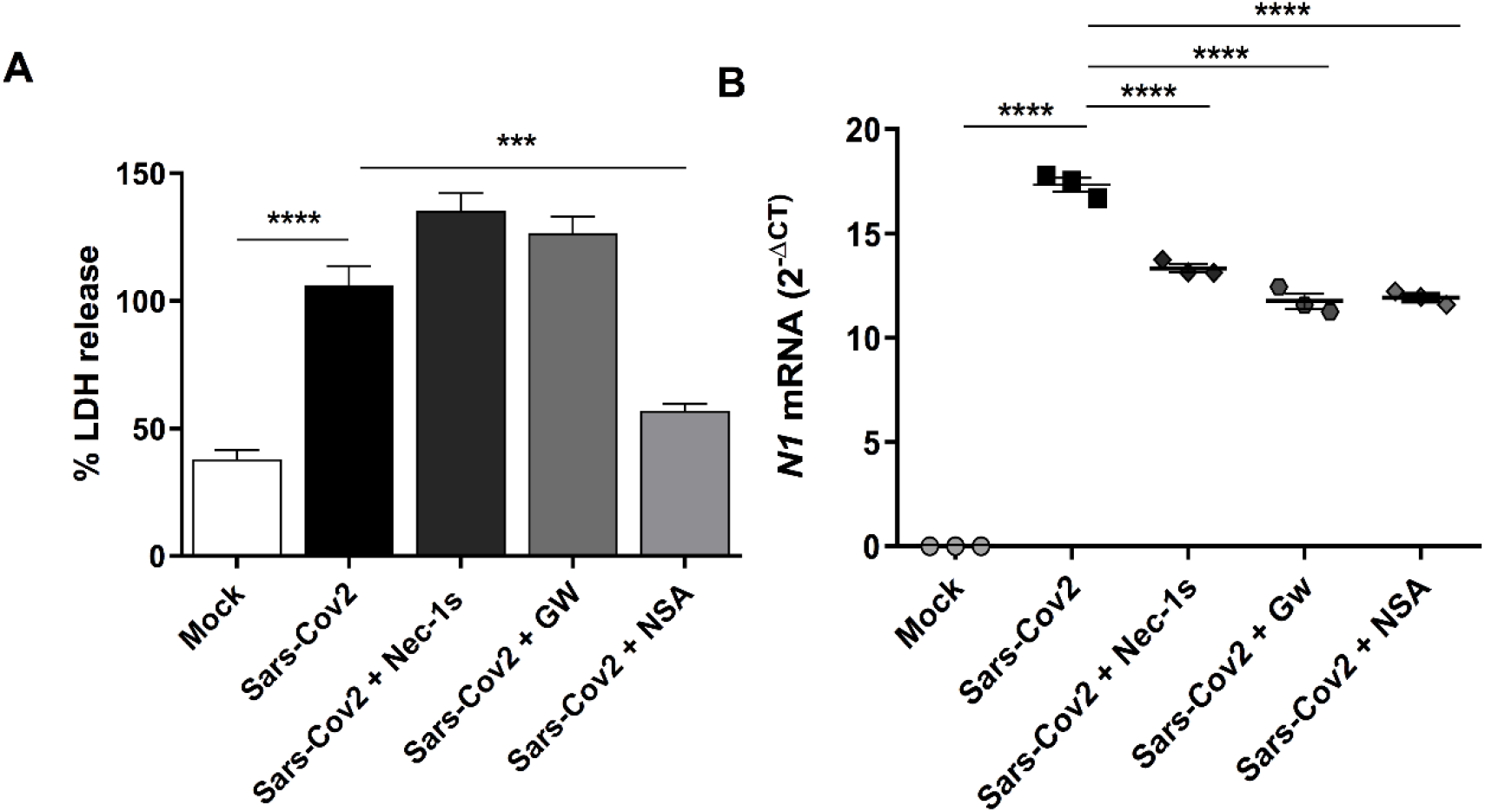
SARS-CoV-2 promotes human monocyte necroptosis. A) Human monocytes (3×10^5^/well) were treated with NEC-1s (30 μM), GW42X (2 μM), or NSA (2 μM) for 1 h prior to SARS-CoV-2 (MOI 0.5) infection for 24 h. Cell death was assessed by LDH release in cell supernatants and expressed as % LDH release. B) Human monocytes (3×10^5^/well) were treated with NEC-1s (30 μM), GW42X (2 μM), or NSA (2 μM) for 1 h prior to SARS-CoV-2 (MOI 0.5) infection for 24 h. RNA was harvested, and SARS-CoV-2 N1 mRNA expression was quantified by real-time PCR using the 2^−ΔCt^ method. Data are representative of at least two independent experiments performed in triplicate and expressed as mean ± SEM. Data were analyzed with one-way ANOVA with Tukey’s post hoc test. *** p≤0.001; **** p≤0.0001.

### SARS-CoV-2–IgG immune complex triggers RIPK3- and MLKL-mediated necroptosis in human monocytes dependently of Fcγ receptors

It has been previously reported that high titers of anti-SARS-CoV-2 IgG promote inflammation by alveolar macrophages [8]. Therefore, we were interested in understanding whether SARS-CoV-2–IgG immune complexes would trigger monocyte necroptosis. We then collected blood from hospitalized mild COVID-19 patients and purified their serum. The clinical characteristics of patients are found in supplementary Table 1. We first measured the concentrations of anti-spike IgG in the serum of COVID-19 patients. We observed two distinct groups of patients regarding their levels of anti-spike IgG – one group presented low levels of IgG (< 500 BAU/mL) while the other one exhibited high levels of IgG (> 500 BAU/mL) (Fig. 2A). For the subsequent experiments, we used the patients’ sera with high concentrations of anti-spike IgG. As a control for the COVID-19 serum, we used serum obtained from healthy volunteer donors before the SARS-CoV-2 pandemic (pre-pandemic serum). Our results demonstrate that incubating human monocytes with either UV-SARS-CoV-2, pre-pandemic serum, or COVID-19 serum only did not induce cell death (Fig. 2B). Similarly, the incubation of monocytes with the mixture of pre-pandemic serum and UV-SARS-CoV-2 was not able to kill monocytes (Fig. 2B). However, stimulating the cells with UV-SARS-CoV-2 immune complexes did promote human monocyte necrosis when compared to negative control (Fig. 2B). We confirmed that SARS-CoV-2 immune complexes triggered monocyte death by comparing LDH release from immune complexes-stimulated cells with that from cells incubated with COVID-19 serum only (Fig. 2C). To evaluate the role of necroptosis-associated proteins on SARS-CoV-2 immune complexes-induced monocyte death, cells were pre-treated with RIPK1, RIPK3 and MLKL inhibitors. Pre-treating human monocytes with RIPK1 inhibitor did not affect cell death (Fig. 2D). In contrast, the treatment of cells with RIPK3 or MLKL inhibitor significantly decreased monocyte death elicited by the immune complexes (Fig. 2D), confirming that UV-SARS-CoV-2 immune complex triggers human monocyte necroptosis. Immune complexes activate monocytes by binding to Fcγ receptors and downstream signaling by Syk kinase [9]. To determine the role of Syk kinase on necroptosis promoted by SARS-CoV-2 immune complex, the cells were pre-treated with a Syk kinase selective inhibitor. This pre-treatment significantly reduced human monocyte necroptosis (Fig. 2D), suggesting that Fcγ receptors are required for SARS-CoV-2 immune complex-induced monocyte necroptosis. Finally, we analyzed LDH release levels with respect to the length of hospital stay of COVID-19 patients and we were not able to find a positive correlation between LDH release and the length of hospital stay (Fig.2E). However, there was a positive correlation between LDH release levels and the need for supplemental oxygen therapy measured in days (Fig. 2F). These data suggest that LDH release levels as a marker of cell death positively correlate with disease severity in patients with COVID-19.

**Fig. 2.**
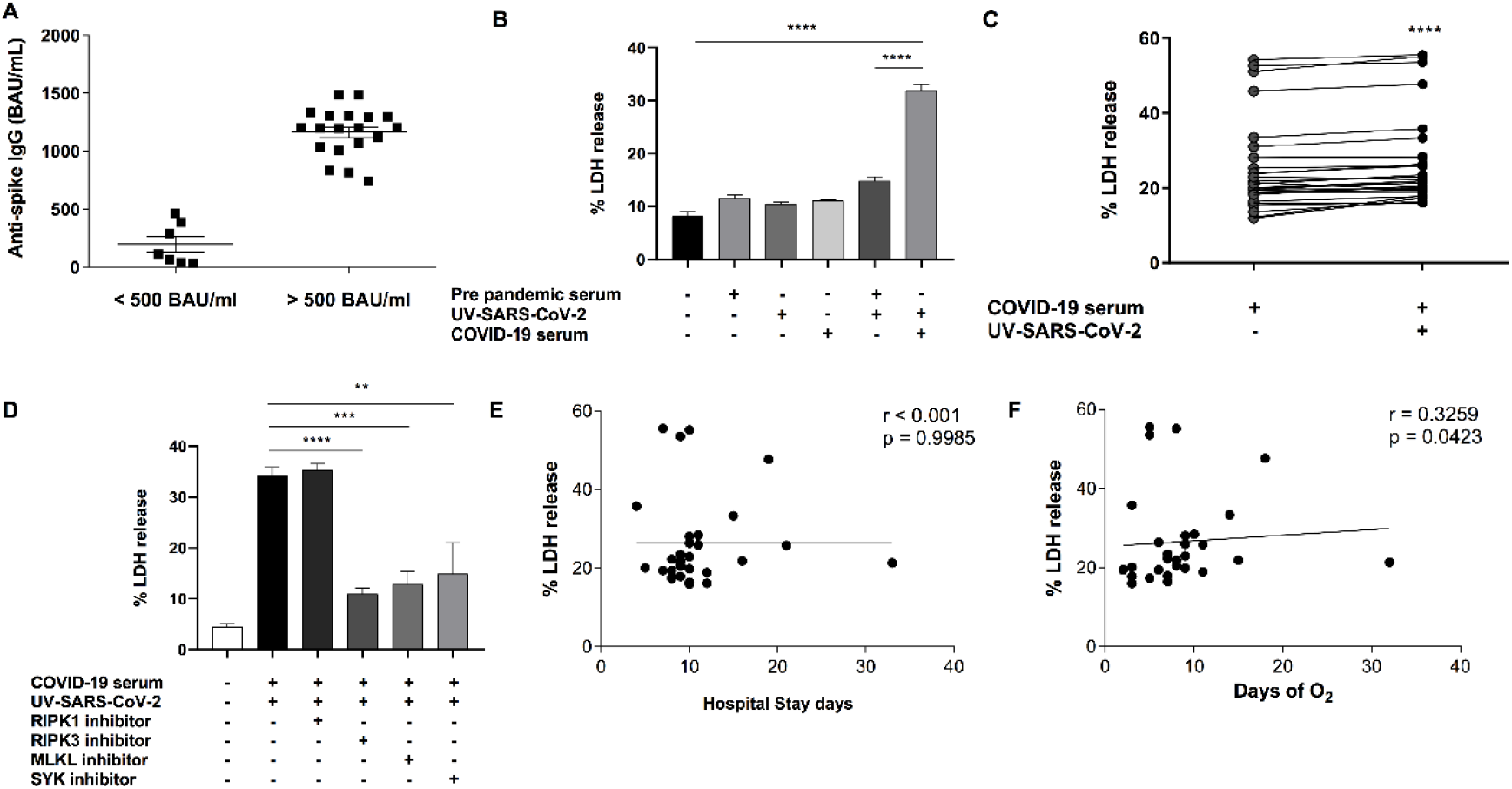
SARS-CoV-2–IgG immune complex triggers RIPK3- and MLKL-mediated necroptosis in human monocytes dependently of Fcγ receptors. A) The concentration of IgG antibodies against the S1 antigen of SARS-CoV-2 was measured in the serum of COVID-19 patients using ELISA and expressed as BAU/mL. B) Human monocytes were stimulated with either UV-SARS-CoV-2, pre-pandemic serum, COVID-19 serum, the mixture of UV-SARS-CoV-2 + pre-pandemic serum, UV-SARS-CoV-2 immune complexes (UV-SARS-CoV-2 + COVID-19 serum), or media only for 24 hours. Afterwards, cell death was assessed by LDH release in cell supernatants and expressed as % LDH release. C) Human monocytes were stimulated with COVID-19 serum only or UV-SARS-CoV-2 immune complexes (UV-SARS-CoV-2 + COVID-19 serum) for 24 hours and LDH release was measured in cell supernatants. D) Monocytes were treated with RIPK1 inhibitor (NEC-1s; 30 μM), RIPK3 inhibitor (GW42X; 2 μM), MLKL inhibitor (NSA; 2 μM), or Syk tyrosine kinase inhibitor (piceatannol; 20 μM) for 1 hour prior to stimulation with UV-SARS-CoV-2 immune complexes (UV-SARS-CoV-2 + COVID-19 serum). Afterwards, cell death was assessed by LDH release in cell supernatants. E) Spearman linear correlation between LDH release levels and length of hospital stay of COVID-19 patients. F) Spearman linear correlation between LDH release levels and time of O_2_ need of COVID-19 patients. Data are representative of at least two independent experiments performed in triplicate and expressed as mean ± SEM. Data were analyzed with one-way ANOVA with Tukey’s post hoc test (B,D) or Wilcoxon matched-pairs signed-rank test (C). ** p≤0.01; *** p≤0.001; **** p≤0.0001.

## Discussion

SARS-CoV-2 has been shown to trigger lytic cell death in different cell types [6, 7, 10]. Therefore, we sought to understand whether SARS-CoV-2 would induce necroptosis in human monocytes. We found that SARS-CoV-2 directly promotes human monocyte necroptosis depending on MLKL activation. Importantly, necroptosis-associated proteins are also involved in SARS-CoV-2 N1 gene expression in monocytes, likely playing a role in SARS-CoV-2 replication.

In COVID-19, antibody-mediated virus neutralization is important for infection control and disease outcomes [11]. However, recent studies suggest that higher anti-SARS-CoV-2 antibody titres correlate with disease severity [12, 13] and promote inflammation by macrophages [7, 8]. We observed two distinct groups of patients regarding their levels of anti-SARS-CoV-2 spike antibodies – one group presenting low levels and the other one exhibiting high levels of antibodies. UV-SARS-CoV-2-IgG immune complexes generated using COVID-19 serum with high levels of anti-SARS-CoV-2 spike antibodies promoted lytic cell death of monocytes. This lytic cell death mode has been confirmed to be necroptosis as the pre-treatment of monocytes with RIPK3 and MLKL inhibitors almost abolished cell death triggered by UV-SARS-CoV-2-IgG immune complexes. IgG immune complexes are recognized by Fcγ receptors. Monocytes express three Fcγ receptors (FcγRI/CD64, FcγRII/CD32 and FcγRIIIa/CD16), which signal through Syk tyrosine kinase [9, 14]. Pre-treating monocytes with a selective Syk tyrosine kinase inhibitor strongly decreased necroptosis induced by SARS-CoV-2 immune complexes, suggesting that the activation of Fcγ receptors may trigger monocyte necroptosis during COVID-19. Necroptotic cells release cytokines, chemokines and other alarmins, which can amplify systemic inflammation and contribute to COVID-19 pathogenesis.

Elevated LDH levels have been used as a marker of severity and a predictor of mortality in patients with COVID-19 [4, 15]. We measured disease severity in our cohort of patients with COVID-19 based on the length of hospital stay and the duration of supplemental oxygen therapy. We were not able to find a positive correlation between the levels of LDH release and the length of hospital stay; however, we did find a significant positive association between the levels of LDH release and the need for supplemental oxygen. These data suggest that elevated LDH levels positively correlate with COVID-19 severity.

In conclusion, our data demonstrate that SARS-CoV-2 itself and SARS-CoV-2 immune complexes trigger human monocyte necroptosis; the latter being mediated by Fcγ receptors. We also provide evidence that elevated LDH levels as a marker of necroptotic cell death are associated with COVID-19 severity.

## Acknowledgments

The authors thank Eduardo Pedrazza for the technical assistance.

---

**Supplementary Table 1.** Clinical characteristics of the study population.

| **Table 1. Clinical characteristics of the study population** |  |
| --- | --- |
| **Variables** | **Values** |
| Gender, n (%) |  |
| Male | 16 (55%) |
| Female | 13 (45%) |
| Age, y (mean ± SD) | 52.896 ± 14.099 |
| Length of hospital stay, days (mean ± SD) | 11,0344 ± 5,480 |
| O<sub>2</sub> need, days (mean ± SD) | 8,379 ± 5,821 |
| SpO<sub>2</sub>, %, (mean ± SD) | 94,965 ± 1,809 |
| BMI, kg/m<sup>2</sup> (mean ± SD) | 28,780 ± 6,450 |
| Definition of abbreviations: SD, standard deviation; Kg, kilograms; y, years. |  |

## References

1. Knoll, R., J.L. Schultze, and J. Schulte-Schrepping, Monocytes and Macrophages in COVID-19. Front Immunol, 2021. 12: p. 720109.

2. Meidaninikjeh, S., et al., Monocytes and macrophages in COVID-19: Friends and foes. Life Sci, 2021. 269: p. 119010.

3. Frank, D. and J.E. Vince, Pyroptosis versus necroptosis: similarities, differences, and crosstalk. Cell Death Differ, 2019. 26(1): p. 99–114.

4. Wu, C., et al., Risk Factors Associated With Acute Respiratory Distress Syndrome and Death in Patients With Coronavirus Disease 2019 Pneumonia in Wuhan, China. JAMA Intern Med, 2020. 180(7): p. 934–943.

5. Karki, R., et al., Synergism of TNF-alpha and IFN-gamma Triggers Inflammatory Cell Death, Tissue Damage, and Mortality in SARS-CoV-2 Infection and Cytokine Shock Syndromes. Cell, 2021. 184(1): p. 149–168 e17.

6. Ferreira, A.C., et al., Correction: SARS-CoV-2 engages inflammasome and pyroptosis in human primary monocytes. Cell Death Discov, 2021. 7(1): p. 116.

7. Junqueira, C., et al., FcgammaR-mediated SARS-CoV-2 infection of monocytes activates inflammation. Nature, 2022.

8. Hoepel, W., et al., High titers and low fucosylation of early human anti-SARS-CoV-2 IgG promote inflammation by alveolar macrophages. Sci Transl Med, 2021. 13(596).

9. Crowley, M.T., et al., A critical role for Syk in signal transduction and phagocytosis mediated by Fcgamma receptors on macrophages. J Exp Med, 1997. 186(7): p. 1027–39.

10. Li, S., et al., SARS-CoV-2 triggers inflammatory responses and cell death through caspase-8 activation. Signal Transduct Target Ther, 2020. 5(1): p. 235.

11. Zohar, T. and G. Alter, Dissecting antibody-mediated protection against SARS-CoV-2. Nat Rev Immunol, 2020. 20(7): p. 392–394.

12. Garcia-Beltran, W.F., et al., Multiple SARS-CoV-2 variants escape neutralization by vaccine-induced humoral immunity. Cell, 2021. 184(9): p. 2523.

13. Li, K., et al., Dynamic changes in anti-SARS-CoV-2 antibodies during SARS-CoV-2 infection and recovery from COVID-19. Nat Commun, 2020. 11(1): p. 6044.

14. Bruhns, P. and F. Jonsson, Mouse and human FcR effector functions. Immunol Rev, 2015. 268(1): p. 25–51.

15. Huang, Y., et al., Serum Lactate Dehydrogenase Level as a Prognostic Factor for COVID-19: A Retrospective Study Based on a Large Sample Size. Front Med (Lausanne), 2021. 8: p. 671667.

