## Supplemental Table 1 for "SARS-CoV-2 immune complex triggers human monocyte necroptosis"

**Table 1. Clinical characteristics of the study population**

| Variables | Values |
|---|---|
| Gender, n (%) | |
| Male | 16 (55%) |
| Female | 13 (45%) |
| Age, y (mean ± SD) | 52.896 ± 14.099 |
| Length of hospital stay, days (mean ± SD) | 11,0344 ± 5,480 |
| $O_2$ need, days (mean ± SD) | 8,379 ± 5,821 |
| $SpO_2$, %, (mean ± SD) | 94,965 ± 1,809 |
| BMI, $kg/m^2$ (mean ± SD) | 28,780 ± 6,450 |

Definition of abbreviations: SD, standard deviation; Kg, kilograms; y, years.
